# Structural features of the fly chromatin colors revealed by automatic three-dimensional modeling

**DOI:** 10.1101/036764

**Authors:** François Serra, Davide Baù, Guillaume Filion, Marc A. Marti-Renom

## Abstract

**Background:** The sequence of a genome is insufficient to understand all genomic processes carried out in the cell nucleus. To achieve this, the knowledge of its threedimensional architecture is necessary. Advances in genomic technologies and the development of new analytical methods, such as Chromosome Conformation Capture (3C) and its derivatives, provide unprecedented insights on the spatial organization of genomes. However, inferring structures from raw contact data is a tedious process for shortage of available tools.

**Results:** Here we present TADbit, a computational framework to analyze and model the chromatin fiber in three dimensions. To illustrate the use of TADbit, we automatically modeled 50 genomic domains from the fly genome revealing differential structural features of the previously defined chromatin colors, establishing a link between the conformation of the genome and the local chromatin composition.

**Conclusions:** TADbit provides three-dimensional built from 3C-based experiments, which are ready for visualization and for characterizing their relation to gene expression and epigenetic states. TADbit is open-source and available for download from http://www.3DGenomes.org.

## Background

Metazoan genomes are organized within the cell nucleus. At the highest level, chromosomes occupy characteristic nuclear areas or “chromosome territories”, separated by inter-chromatin compartments [1]. Chromosomes undergo additional levels of arrangements and organize themselves into the so-called A and B compartments [2], which in turn are composed of Topologically Associating Domains (TADs), defined as regions of the DNA with a high frequency of self-interactions [3-5]. Determining the three-dimensional (3D) organization of such genomic domains is essential for characterizing how genes and their regulatory elements arrange in space to carry out their functions [6]. Chromosome Conformation Capture (3C) [7] and its derived methods (here referred to as 3C-based methods) are now widely used to elucidate the spatial arrangement of genomes [8]. Although the frequency of interactions between loci can be used as a proxy for their spatial proximity, 3C-based contact maps do not easily convey all the information about the spatial organization of a chromosome. This information, however, can be inferred using computational methods [9]. Here we present TADbit, a Python library for the analysis and modeling of 3C-based data. TADbit takes as input the sequencing reads of 3C-based experiments and performs the following main tasks: (i) pre-process the reads, (ii) map the reads to a reference genome, (iii) filter and normalize the interaction data, (iv) analyze the resulting interaction matrices, (v) build 3D models of selected genomic domains, and (vi) analyze the resulting models to characterize their structural properties (Fig. 1). TADbit builds on existing partial implementations of methods for 3D genomic reconstruction [10-20]. As a validation of the model-building module of TADbit, a systematic analysis of its limitations has shown that 3D reconstruction of genomes based on 3C-based data can produce accurate 3D models [21].

**Figure 1.**
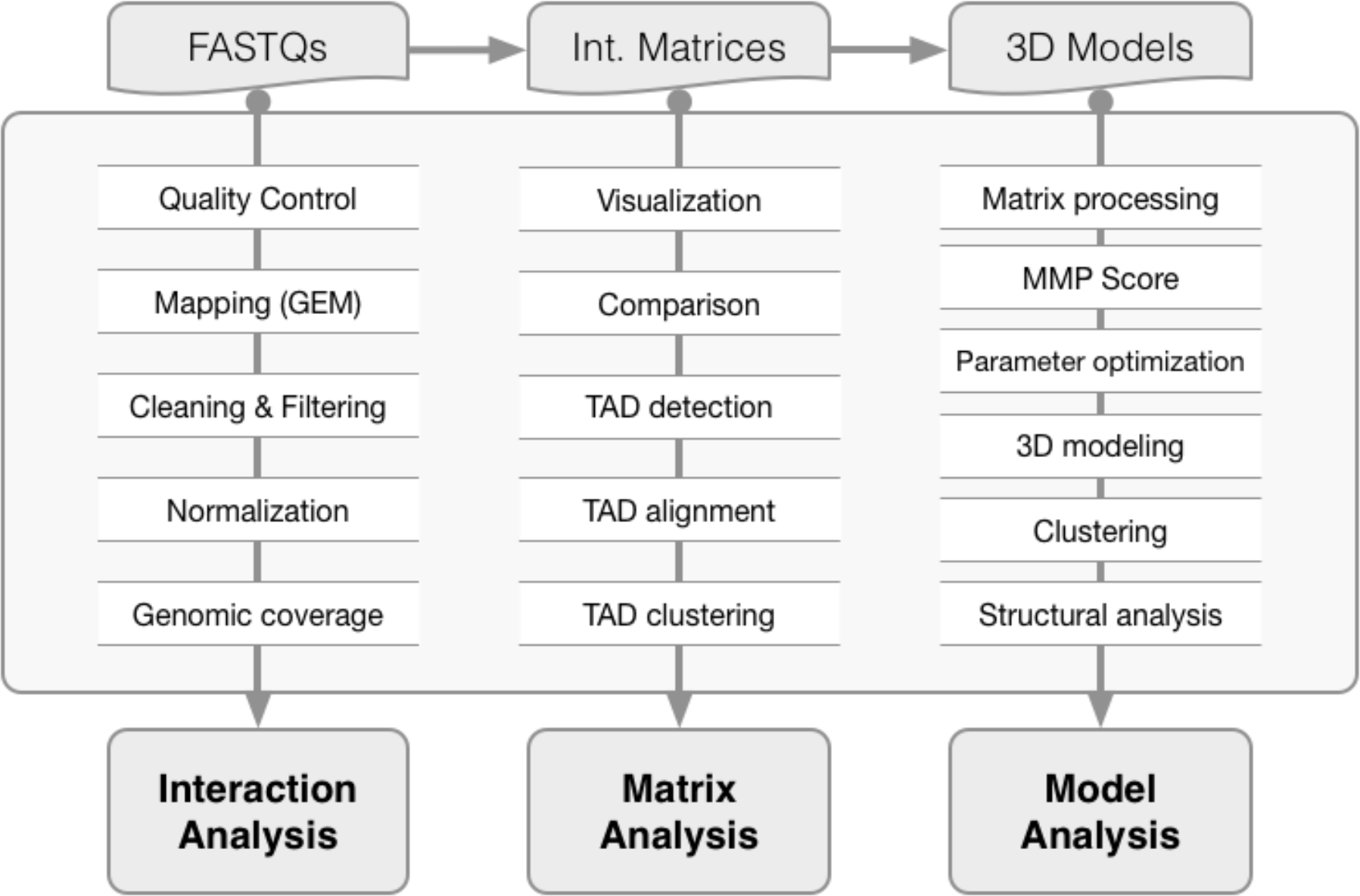
TADbit flowchart. Main functions of the TADbit library from FASTQ files to 3D model analysis. TADbit accepts many input data types such as FASTQ files, interaction matrices and 3D models. A series of python functions in TADbit (**Supplementary Text**) allow for the full analysis of the interaction data, interaction matrices as well as derived 3D models.

Here we used TADbit to model and analyze 50 genomic domains of the *Drosophila melanogaster* genome. It was shown that the *Drosophila* genome consists of five distinct chromatin types determined by mapping 53 broadly selected chromatin proteins and four key histone modifications [22]. The chromatin types were labeled with colors and comprise “blue” chromatin, enriched in Polycomb group proteins and H3K27 methylation, “green” chromatin, bound by HP1 and located at peri-centromeric regions, “yellow” and “red” chromatin, harboring distinct classes of active genes, and “black” chromatin, covering more than 40% of the *Drosophila* genome and characterized by low occupancy of most chromatin markers. More recently, genome-wide 3C-based interaction maps in *Drosophila* revealed that TAD boundaries are gene-dense, highly bound by transcription factors and insulator proteins and correspond to transcribed regions [5, 23]. Moreover, it was shown that the active red and yellow chromatin types preferentially locate at TAD borders while the others preferentially locate inside TADs. This work highlighted the existence of interplay between the structural organization of genomic domains and their chromatin composition. Similar relationships have also been observed in other organisms, including mouse and human [24-27].

To further characterize the structural properties of the *Drosophila* chromatin types, we have used TADbit on available Hi-C data. By building 3D models of genomic domains covering more than 50 Mb of the *Drosophila* genome, we show that the five previously described chromatin colors are characterized by distinct structural properties. Black chromatin is a compact, dense and closed chromatin fiber. In comparison, the heterochromatic types blue and green are more open and accessible. Finally, the yellow and red types feature a loose and open chromatin, potentially accessible to proteins and transcription factors responsible for regulating resident genes.

## Results

### Chromatin interaction maps of the *Drosophila melanogaster* genome

The TADbit pipeline starts from raw data (*i.e*. reads generated from a 3C-based experiment). We downloaded SRA files from the NCBI Gene Expression Omnibus under accession number GSE38468 [23], and converted them to FASTQ files using the SRA Toolkit [28]. The dataset contained three separate Hi-C experiments [2] performed on *Drosophila* Kc167 cells using the restriction endonuclease HindIII, consisting of one biological replicate (SRR398921) and two technical replicates (SRR398318 and SRR398920), labeled here as “BR", “TR1” and “TR2”. They comprised about 194, 67 and 112 million paired-end reads, respectively (Table 1). A quality check of the first million reads in each of the FASTQ file showed that the average PHRED scores [29] were higher than 25 across each of the 2x50 bp paired-end reads, which is indicative of good quality. Moreover, TADbit assessed that more than 95% of the reads had undergone digestion during the Hi-C experiment and only ∽2% of the reads contained dangling ends *sensu stricto* (reads starting with a digested restriction site, Fig. S1). Next, the paired-end reads were aligned in TADbit to the *Drosophila* reference genome (dm3) using the GEM mapper [30] with a previously proposed iterative mapping strategy [31]. With this strategy, 67.0% to 77.8% of the original reads could be uniquely mapped (Table 1). After discarding those with only one mapped end, the number of mapped pairs diminished (50.2% to 63.5% of the original reads). These numbers were similar to those reported in the original experiments [23]. After mapping, the reads were further filtered as previously described [31], resulting in about 48, 24, and 41 million valid pairs (or interactions) for the BR, TR1 and TR2 experiments, respectively (Table 1). Finally, the filtered interaction maps were normalized using the iterative correction and eigenvector decomposition (ICE) procedure [31], also implemented in TADbit (Fig. 2a). The resulting interaction matrices were highly correlated (Fig. 2b,c,d), which prompted us to merge the input reads into a single dataset of more than 372 million reads. The new dataset, referred to as “SUM”, was also automatically filtered and normalized by TADbit (Fig. 2e,f). The interaction map from the SUM dataset shows all the previously described features of the 3D organization of the *Drosophila* genome, including the chromosome arm territories, the clustering of centromeres and the infrequent interactions between telomeres.

**Table 1.**
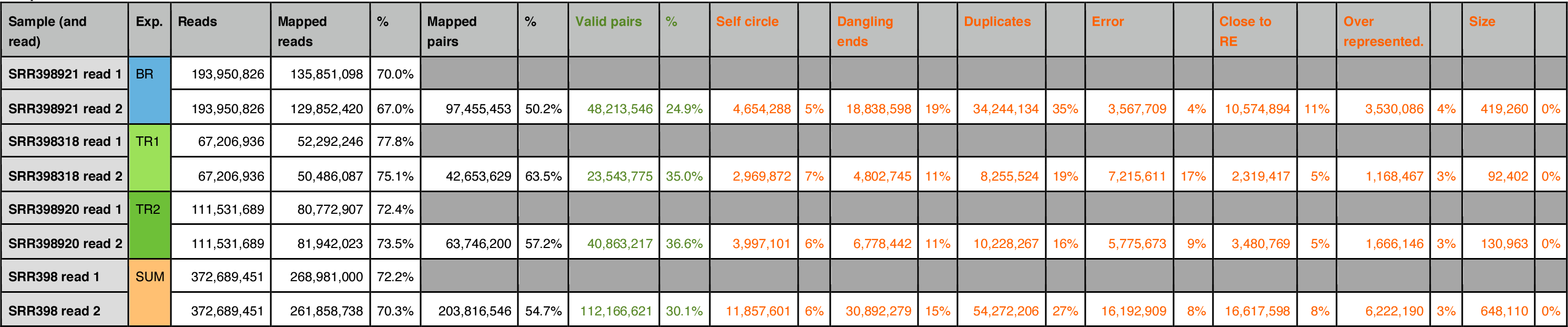
TADbit mapping and filtering of the Hi-C experimental results. Columns indicate the number of reads per experiment (“Reads”), those that were uniquely mapped (“Mapped reads”), pairs of reads mapped in both ends (“Mapped pairs”) and finally the number of reads that were remained after filtering (“Valid pairs”). The filtered reads were quantified by “Self circles”, “Dangling ends”, “Duplicates”, “Close to RE”, “Over represented", and “Size" filters.

**Figure 2.**
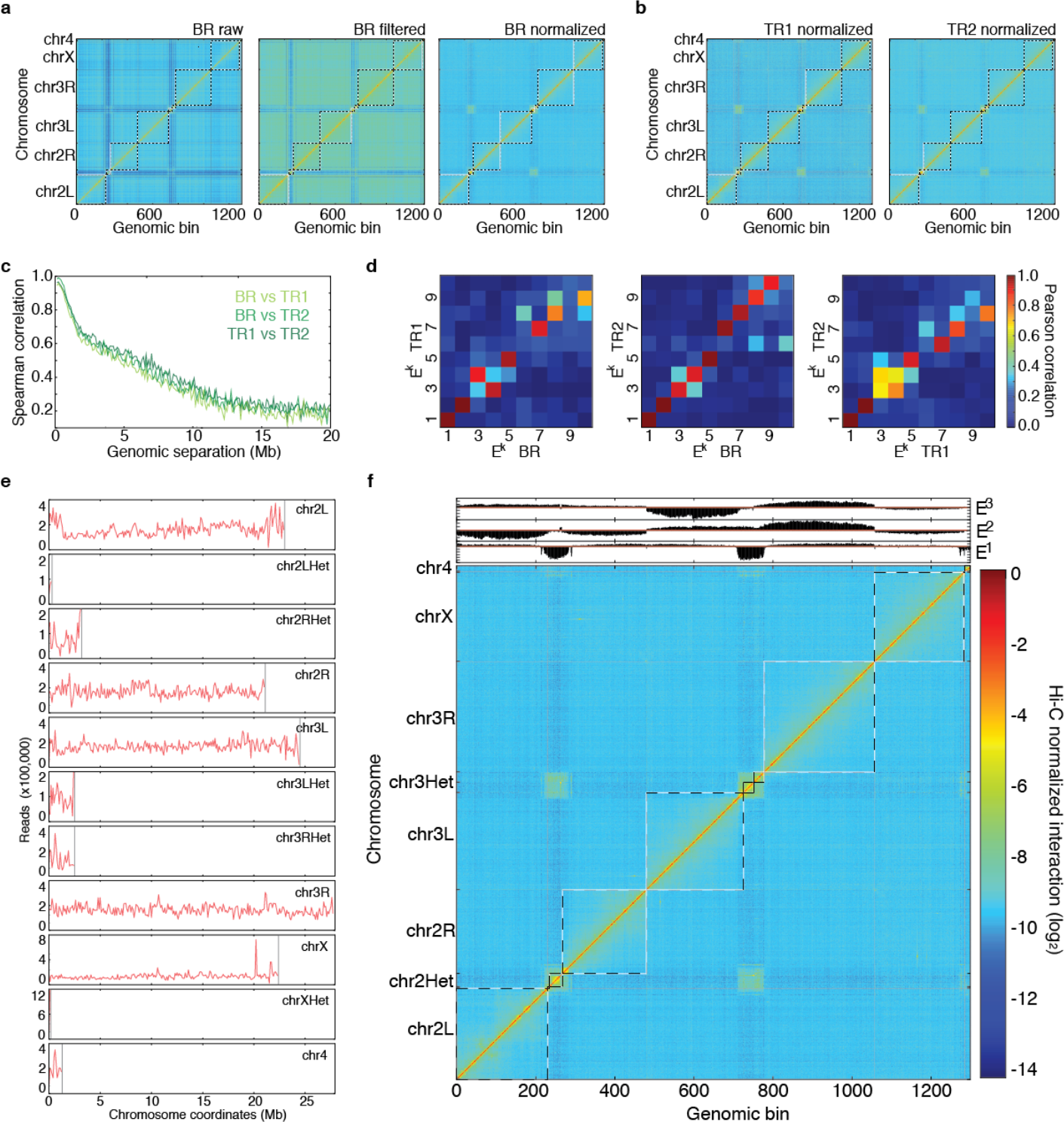
Hi-C interaction maps at 100 kb resolution for the entire *Drosophila* genome. (a) Raw, filtered and normalized genome-wide interaction maps for the BR dataset. Only after the normalization of the data, the enriched interaction between centromere regions of the Drosophila chromosomes can be observed. (b) Normalized maps for the TR1 and TR2 datasets. (c) Comparison of the normalized Hi-C maps between the three datasets at 100 kb resolution. The Spearman correlation was computed between off-diagonal regions as a function of their genomic distance. (d) Matrices of Pearson correlation coefficients of main eigenvectors from the three Hi-C datasets (that is, BR, TR2 and TR2). The data shows the expected high correlation of the top three eigenvectors [31]. (e) Genomic coverage of the mapped reads per chromosome from the SUM dataset. (f) Hi-C normalized interaction matrix at 100 kb resolution for the SUM dataset. The three main eigenvectors of the normalized interaction matrix mark the position of centromeres (E^1^), chromosomes (E^2^), and chromosome arms (E^3^). TADbit automatically generated all the plots in the figure.

### The *Drosophila* genome is partitioned into TADs of different robustness

Next, we generated 10 kb resolution interaction maps of the *Drosophila* genome to which we applied a new TAD boundary detection algorithm implemented in TADbit (Materials and Methods). This algorithm uses a change-point detection approach inspired from methods used to identify copy number variations in CGH experiments [32]. Briefly, we use Poisson regression to find the most likely segmentation of the chromosome in *m* TADs and choose the value of *m* associated with the optimal Bayesian Information Criterion. In addition to the optimality of the solution, the main advantage of the new algorithm is the assignment of a robustness score to each TAD boundary (Materials and Methods). TADbit identified a total of 689 TADs with an average length of 162.8 kb (ranging from 20 kb to 1.5 Mb), representing larger TADs than previously reported [23]. Given the hierarchical organization of the genome [8], we set out to assess whether the difference was due to the identification of new borders or to the merging of the identified TADs. We downloaded the interaction matrices and the TAD borders as defined by Hou *et al*. [23] (referred to as the original definition) and compared them to the borders obtained by running TADbit on these interaction matrices (Fig. 3a,b,c). To this end we used the TADbit module to align multiple TAD boundaries from several experiments (Materials and Methods and Fig. 3d). Overall, 81% of the borders defined by TADbit align within 20 kb of an original border when using the TADbit definition as reference (Fig. 3e). The number decreases to 67% of the borders when using the original definition as a reference. By forcing TADbit to identify the same number of borders as the original definition (1,110 borders), the agreement increases to 74% within 20 kb. For comparison, the agreement of the TADbit border definitions between the three independent Hi-C experiments (BR, TR1 and TR2) is about 90%. The degree of similarity between the original and the TADbit definitions points to a variation of the algorithm sensitivity more than to real discrepancies (see Fig. 3d for instance). Moreover, the borders present only in the TADbit definition usually have a weak strength. Indeed, the agreement increases to 94% by comparing borders of 6 or higher strength as defined by TADbit. In summary, our results confirm the previously described TAD level partitioning of the *Drosophila* genome and provide a new algorithm to identify TAD borders and to assign a strength score to them. Such strength score could later be used to characterize the hierarchical organization of the genome in TADs or as an indicator of the confidence in the prediction (Fig. S2).

**Figure 3.**
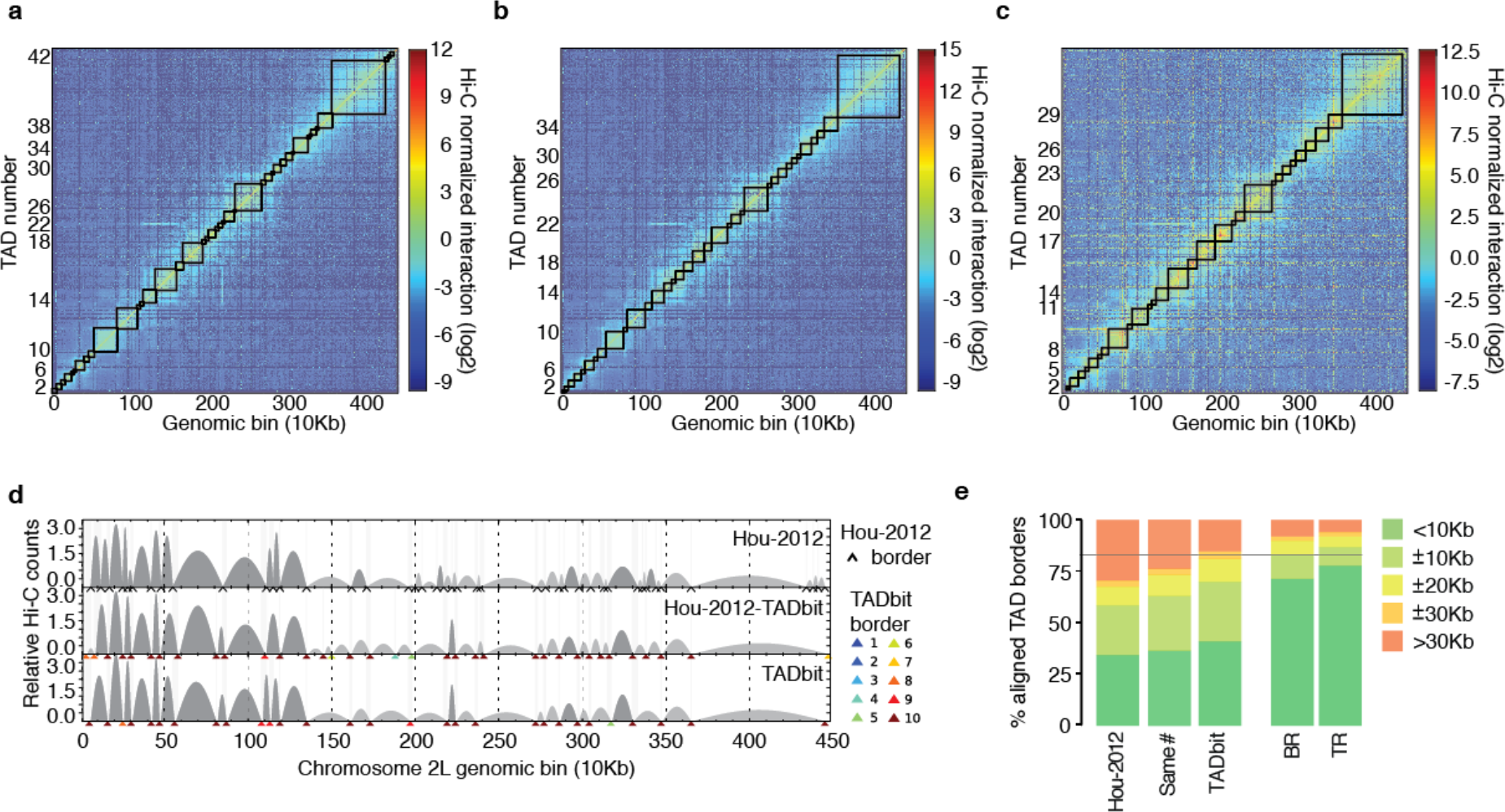
TAD border detection and comparison with the results from Hou *et al*. [23] (a) Hi-C normalized interaction matrix at 10 kb resolution for the first 4.5 Mb of chromosome 2L in the *Drosophila* genome. Interactions matrix and TAD borders were obtained from published data [23]. (b) Hi-C normalized interaction matrix from the same genomic region and resolution as in panel a. The interaction counts are as previously published [23] but the TAD borders are those defined by TADbit. (c) Hi-C normalized interaction matrix from the same genomic region and resolution as in panel a. Interaction data and TAD borders are both generated by TADbit. (d) TAD border alignments between the three differently processed experimental data: borders defined in Hou *et al*. [23] (Hou-2012, top graph), borders defined by TADbit using the Hou-2012 matrix (mid graph), and borders and matrix determined by TADbit (bottom graph). Dark and light grey arches indicate TADs with higher and lower than expected intra-TAD interactions, respectively. TAD borders are indicated with a black arrow for the Hou-2012 defined borders and by color arrows for the TADbit identified borders. TADbit border robustness (from 1 to 10) is identified by a color gradient from blue to red. (e) Comparison of the agreement between the aligned TAD borders in the three datasets. As a reference, the horizontal grey line indicates a ±20 kb (2 bins) agreement between the biological replica (BR) and the first technical replicate (TR1) as determined by TADbit. The plots in panels *a* to *d* were automatically generated by TADbit.

### Automatic modeling of 50 genomic regions of the *Drosophila* genome

Next, we used TADbit to model the 3D structure of 50 selected genomic regions of about 1 Mb each (Table S1). The 50 regions were selected based on their chromatin colors composition [22]. The selection included the top ten regions of the genome most enriched in each of the five defined chromatin colors. Given the nonhomogenous distribution of chromatin colors in the *Drosophila* genome, where the genome is composed of large stretches of black chromatin interspersed by shorter domains of blue, yellow and red chromatin (green chromatin is an exception, as it is mainly found in peri-centromeric regions and on chromosome 4), finding continuous 1 Mb stretches of chromatin for the blue, yellow and red colors was not always possible (Fig. 4a). For instance, the highest red coverage in a 1 Mb region of the genome was only 22%. For yellow and blue, the maximum coverage was 48% and 52%, respectively, whereas for black and green chromatin types the maximum coverage was 98% and 100%, respectively.

**Figure 4.**
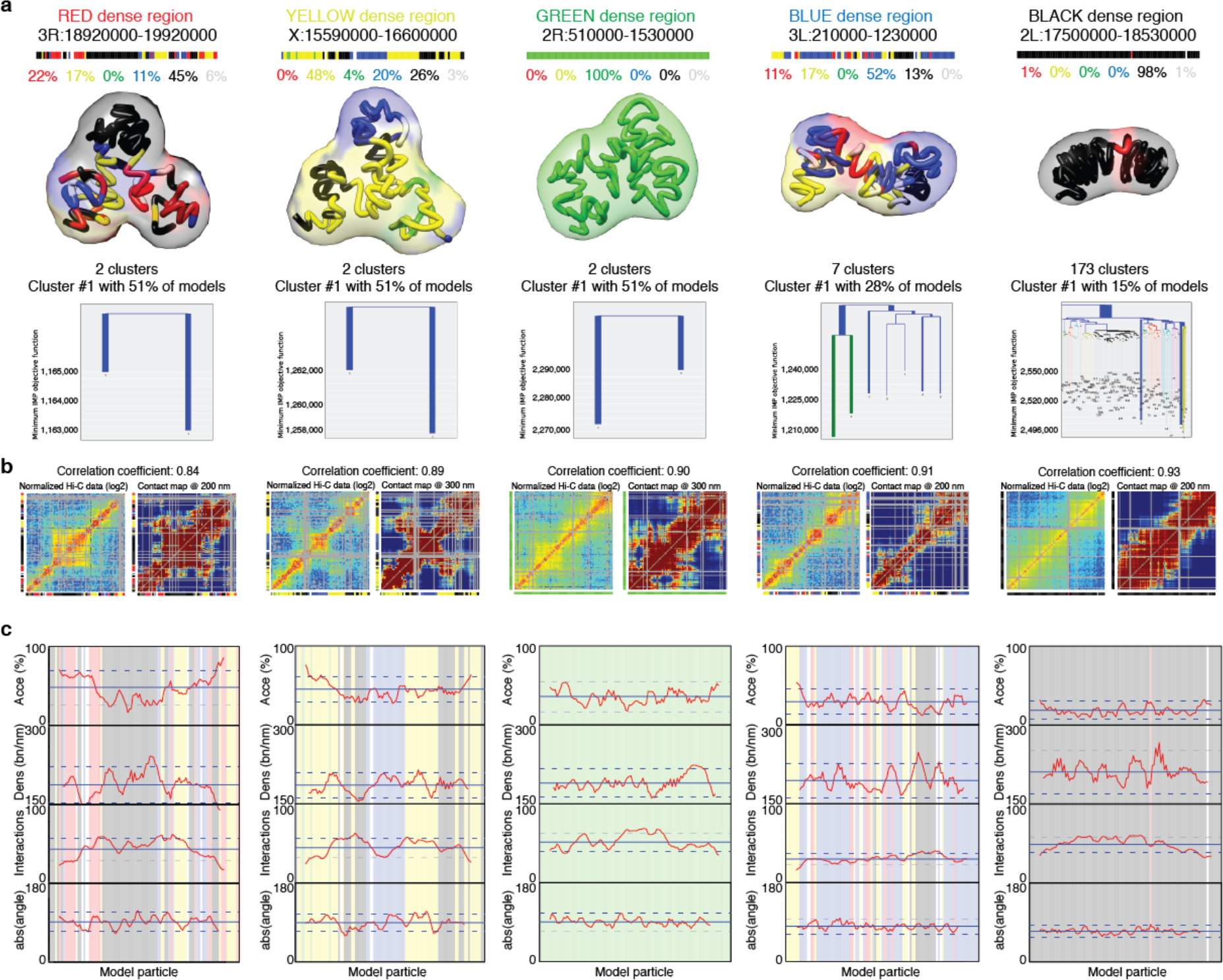
TADbit 3D models and structural properties. (a) Genomic coordinates, chromatin color proportions, 3D models and structural clustering for the five regions with highest coverage for each color in the *Drosophila* genome. The ensemble of models for cluster number 1 (the most populated cluster) for each color is represented by its centroid as a solid tube colored by its particle colors. The ensemble around the centroid is simulated by a transparent surface covering a Gaussian smooth surface 150 nm away from the centroid. Figures of 3D models were produced by Chimera [41]. The structural clustering of the 2,000 models produced per region were aligned with TADbit and clustered by structural similarity. Most modeled regions segregate into two large clusters corresponding to mirror images of each other. (b) Comparison of the input interaction Hi-C matrix to a contact map from the 2,000 built models per region, with Spearman correlation coefficient. (c) Structural properties by particle are shown for accessibility (percentage), density (bp per nanometer), interactions (number), and angle (degree). The background of the plot represents the color assigned to each of the particles in the models. TADbit automatically generated all plots.

All the selected genomic domains yielded a Matrix Modeling Potential (MMP) score [21] ranging from 0.85 to 0.96, which is predictive of high accuracy models (Table S1). To model the 3D structure of the 50 regions, we used as input the Hi-C interaction matrix where each 10 kb bin was represented as a spherical particle in the model. All the particles were restrained in space based solely on their measured interactions, chain connectivity and excluded volume. Finally, the modeling parameters were optimized by maximizing the correlation between the contact map of the models and the input Hi-C interaction matrix (Materials and Methods and Table S1). All the 50 modeling exercises resulted in high correlations between the contact maps and the Hi-C interaction matrices, ranging form 0.83 to 0.93 (Fig. 4b and Table S1). All together, the modeled regions covered a total of 51.8 Mb of the *Drosophila* genome, forming the largest dataset of genomic regions modeled at 10 kb resolution (Fig. S3).

### Structural properties of the *Drosophila* chromatin colors

The generated models were automatically analyzed by TADbit to further characterize their structural properties. In particular, among the set of descriptive measures available in TADbit, we calculated four main structural properties for each particle (genomic bin) in the models (Materials and Methods). Those included: (i) *accessibility*, measuring how accessible from the outside a particle is; (ii) *density*, measuring the local compactness of the chromatin fiber; (iii) *interactions*, counting the number of particles within a given spatial distance from a selected particle; and (iv) *angle*, measuring the angle formed by a particle and its two immediate neighbor particles. To assess whether the different occupancy of proteins and chromatin modifications defining the five colors of chromatin had an influence on the 3D structure of the genome, we assigned to each particle one of the five chromatin colors if at least 50% of the 10 kb region was covered by this chromatin type [22]. Particles with non-homogenous colors were assigned to the undefined “white” color. These four measures provided an overview of the structural properties of each color in a particle-based manner. Models with decreasing amount of black, blue and green particles resulted in less compact and regular structures compared to those enriched in blue or black particles (Fig. 4c). For example, the top black region (98% black, 1% red and 1% white) had low accessibility throughout, combined with a relatively high density (interestingly, the lowest density for that region corresponds to the only red particle), high number of interactions and closed angle between particles (Fig. 4c last column).

Overall, the chromatin colors showed distinct structural properties (Fig. 5). For example, black chromatin was the least accessible (median accessibility 26.5%), compared to green and blue (median accessibilities 34.4% and 34.3%, respectively) and to yellow and red (median accessibilities 46.5% and 51.6%, respectively). Black chromatin also featured the highest density in our models (median 212 bp/nm). This was slightly more than blue (207 bp/nm) and substantially more than green, yellow and red (182 bp/nm, 180 bp/nm, and 179 bp/nm, respectively). The chromatin type with most interactions was green (median 48.7 interacting particles within 250 nm) followed by black (45.3), yellow (43.7), blue (41.9), and red (37.9) chromatin. Finally, yellow and red chromatin featured the most extended fibers (median absolute angles 94.6° and 89.7°, respectively), compared to blue (85.3°), green (82.6°) and black (80.3°). Taken together, the 3D models generated by TADbit indicate that the chromatin types of *Drosophila* have intrinsic and distinctive structural properties.

**Figure 5.**
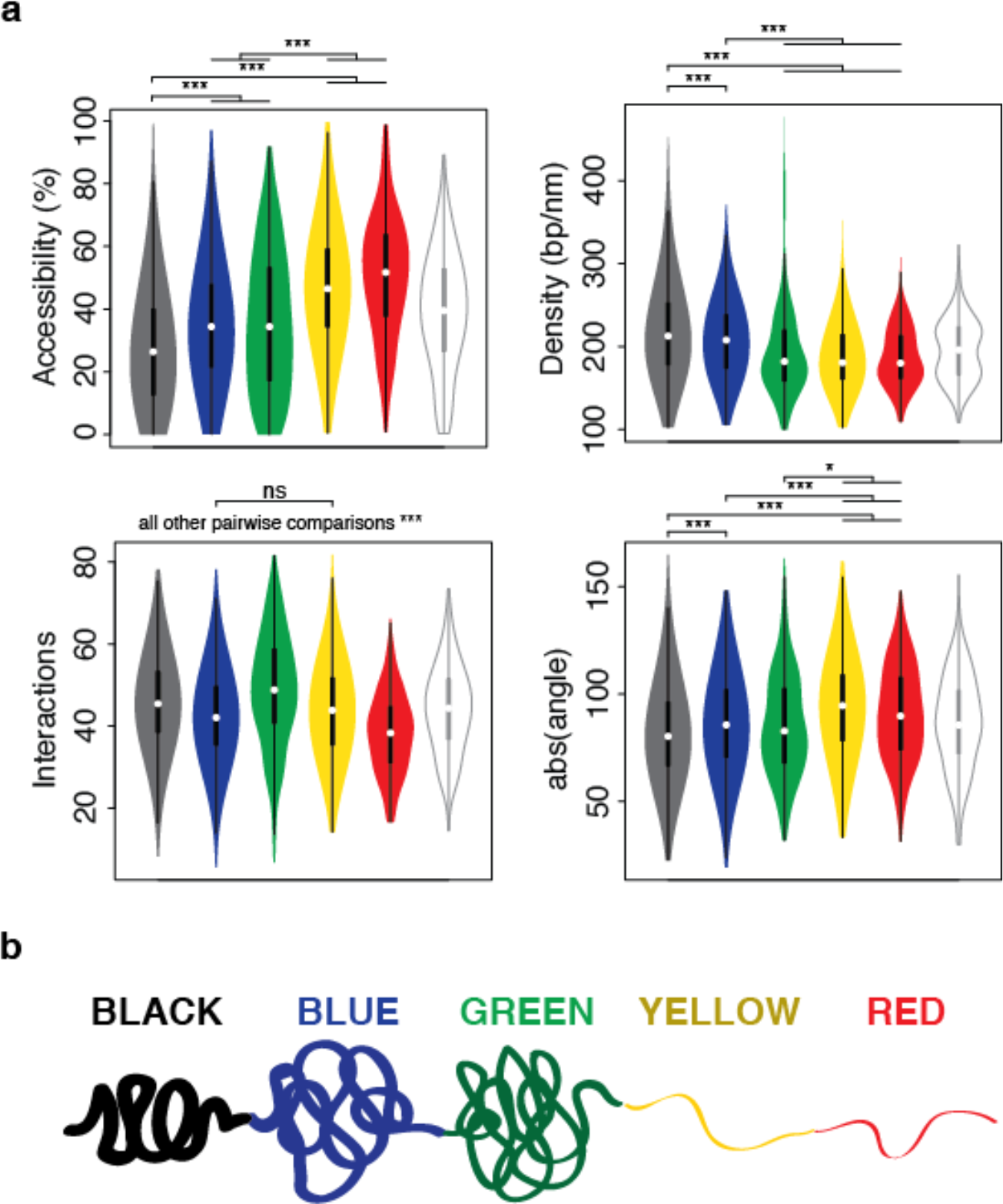
Structural properties of the five described chromatin colors. (a) Distribution of each of the four structural properties (that is, accessibility, density, interactions, and angle) grouped by chromatin colors (including the undefined “white” color for particles of non-homogeneous coloring). Statistical significance of the differences as computed by Tukey’s ‘Honest Significant Difference’ test (*: p < 0.01, ***: p < 0.001, ns: non-significant). (b) Schematic representation of the structural properties of the five colors for the *Drosophila* chromatin.

## Discussion

Here we introduced the first comprehensive computational library for 3C-based data handling, 3D modeling and model analysis. TADbit’s main scopes are: (i) read quality control and design of the mapping strategy; (ii) mapping of reads to the reference genome; (iii) interaction map filtering and normalization; (iv) interaction matrix analysis, including matrix comparison, TAD detection and TAD alignment; (v) 3D modeling of genomes and genomic domains; and (vi) 3D model analysis. A complete list of the computational functions implemented in TADbit is provided as Supplementary Text, each of which is more deeply described in the TADbit online documentation (http://3dgenomes.github.io/TADbit) together with complete tutorials covering each step from sequencing data to 3D model analysis.

Here, TADbit was used to map, normalize, model and analyze a previously published dataset of Hi-C experiments in *Drosophila* Kc167 cells [23]. The resulting interaction maps confirmed previously described features of the 3D organization of the *Drosophila* genome, including the chromosomal arms territories, the clustering of centromeres, the separation of telomeres and the partitioning of the genome into TADs. The mapping strategy using GEM [30] gives similar results to those using Bowtie [33] and alternative protocols [23]. The TAD detection module of TADbit identified less TADs than previously reported. Part of the discrepancy is due to the arbitrary definition of TADs and the lack of knowledge about their real structure. TADbit contributes a scoring of TAD borders, which hints to the hierarchical nature of the genome organization in 3D. Indeed, the current debate on the exact nature of TADs and the existence of a hierarchical organization of domains at different resolutions [25] could be aided by assigning a score of robustness to each of the detected domain boundaries.

We also used TADbit to automatically model a total of 50 genomic regions across the *Drosophila* genome, covering nearly 52 Mb at a resolution of 10 kb, which represents the larges dataset of 3D structures of genomic domains available today. TADbit can also be used to produce genome-wide models. Unfortunately, the computational time required to model the entire Drosophila genome at 10 kb resolution is very high and thus would not be available to many research groups world-wide. Our 3D models reveal distinct structural properties for the previously identified chromatin colors in *Drosophila*. It has been shown that the five types of *Drosophila* chromatin not only differ in protein composition but also in biochemical properties, transcriptional activity, histone modifications, replication timing, and DNA binding factors targeting. They also differ in the sequence properties and the functions of the embedded genes. Now we demonstrate that the chromatin types also have specific and distinctive structural features (Fig. 5b). Importantly, these results shed light on the nature of the elusive black chromatin. Most chromatin markers are depleted in this environment, including those responsible for active repression of transcription. It is thus unclear how genes are maintained silent and why transcription factors do not bind to their consensus sequence in black chromatin. Our results suggest that part of the answer is that black chromatin is very compact and inaccessible to external factors. The high curvature of black chromatin fibers in the models suggests that those regions are intrinsically ordered or that they are compressed. The enrichment of the linker histone H1 in black chromatin may account for all these properties.

The previous conception of heterochromatin was closer to green (HP1-bound) or blue (Polycomb-bound) chromatin types. Interestingly, both of them are more accessible than black chromatin, yet green chromatin has a higher number of interactions. This indicates that green chromatin, compared to black chromatin, is a more open but irregular structure where specific interactions are more plausible within a distance cut-off. In contrast, the closed and regular organization of black chromatin results in fewer likely unspecific interactions per particle. This may somehow be related to the observation that the expression of some genes translocated to HP1-bound regions tends to fluctuate, a phenomenon known as position effect variegation [34]. We speculate that genes caught in this chromatin environment may be trapped in the local entanglement and physically locked away from their enhancers.

Both yellow and red chromatin exhibit the most different structural features compared with black chromatin. Their 3D models are open and accessible, which is consistent with the fact that those regions are mostly transcribed and bound by many transcription factors. However, the overall protein occupancy in red chromatin is substantially higher than in yellow chromatin, yet their overall structural properties are relatively similar. This suggests that the extraordinary occupancy observed in red chromatin is not necessarily rooted in its conformational properties, but rather in mechanisms that operate at a finer scale.

Additional studies will be needed to further investigate the molecular mechanisms associated to the structural properties of the chromatin types. However, our 3D models, as well as their correlation with the epigenetic features, are a firm basis for future investigation on chromatin occupancy by proteins and it spatial organization.

## Materials and Methods

### FASTQ quality check

The TADbit pipeline starts by performing a quality control on the raw data in FASTQ format. This quality check is similar to the tests performed by the *FastQC* program [35] with adaptions for Hi-C datasets (see Fig. 1 and Fig. S1). In particular, it averages the PHRED scores along the sequenced reads as well as the proportion of “N” at each position. Additionally, the distribution of three categories of sequence sites can be plotted: (i) the proportion of undigested restriction enzyme (RE) sites; (ii) the proportion of ligated sites; and (iii) the proportion of non-ligated digested sites *(i.e*., the rest of sites). These three categories might be informative about possible artifacts in the 3C-based experiments. The first and the second categories are expected to be constant along the sequenced reads, although the proportion of ligation sites detected towards the read ends might be lower as a consequence of the interference of the streptavidin beads with the ligation of the sequencing adapters (as observed in the samples Hou-2012). The third category of sites consists of dangling-ends *sensu stricto (i.e*., the reads starting with a digested restriction site). These statistics are useful to *a priori* assess the efficiency of the digestion (by comparing the proportion of ligated and digested sites versus undigested sites) as well as of the efficiency of ligation (by comparing the proportion of dangling ends versus ligated sites). For example, in the case of the samples Hou-2012, the digestion and ligation efficiencies are very similar between TR1 and TR2, while BR results in higher ligation efficiency.

### Iterative mapping

TADbit implements an iterative mapping strategy that is a slightly modified version of the original ICE method developed for the HiClib library [31]. The minimal differences with the original ICE method are the mapper used (TADbit uses GEM [30]) and a more flexible way to define the position of the iterative mapping windows, which can now be fully defined by the user.

### Fragment based filtering

The filtering strategy implemented in TADbit builds on previously described protocols [31] to correct all the computationally detectable experimental biases/errors. After mapping, TADbit can filter the reads depending on ten criteria (Fig. S4), which can be applied individually or as a set of filters. These criteria include:

1. *Self-circles*: reads mapped on different strand within a single RE fragment. The first read, in genomic coordinates, needs to be mapped in the reverse strand.
2. *Dangling-ends:* reads mapped on different strand within a single RE fragment. The second read, in genomic coordinates, needs to be mapped in the reverse strand.
3. *Errors:* reads mapped within a single RE fragment, both mapped on the same strand.
4. *Extra dangling-ends:* reads mapped on different RE fragment but that are close enough (< 500 bp) to be likely from one single DNA fragment. Additionally, the reads need to be mapped on different strands with the second read mapping on the reverse strand.
5. *Too close to a RE site:* reads mapped to the start position too close (< 5 bp) to the RE cutting site, these are also referred to as “semi-dangling-ends” as they actually have, strictly speaking, at least one end dangling.
6. *Too short:* reads mapped within a small restriction fragments (< 100 bp). Such RE fragments are smaller than the library read length and are likely an artifact.
7. *Too large:* reads mapped within a large restriction fragments (> 100 kb, P < 10^-5^ to occur in a randomized genome); they likely represent poorly assembled or repeated regions in the reference genome.
8. *Over-represented:* reads coming from the top 0.5% most frequently detected RE fragments. Such reads are likely arising from PCR artifacts or may represent fragile regions of the genome as well as genome assembly errors.
9. *Duplicates:* duplicate removal of read-pairs mapping in the exact same genomic coordinates are removed as probable PCR duplicates.
10. *Random breaks:* reads mapping too far (> 500 bp) from the RE cutting site. This filter excludes potential non-canonical enzyme activity or random physical breakage of the chromatin.

### Interaction matrix cleaning and normalization

Once filtered, the read-pairs are binned at a user-specified resolution (bin size) depending on the matrix density required by the analysis to be performed. However a minimum amount of counts per bin is usually required for the normalization of the data. Indeed, the ICE correction [31], which converges when all the rows of the matrix have an equal number of normalized interactions, would fail when the density of the matrix is too low *(e.g*., rows with more than 50% of cells with no count). By default the ICE implementation excludes from the normalization the rows that represent the lowest 2% of the matrix. However, depending on the dataset, this number can greatly affect the result by either masking valid columns or keeping sparse columns that would impede the convergence of the normalization algorithm. To determine the threshold amount of interactions for masking columns, TADbit proceeds in two steps. First, the columns with zero counts are removed. Second, a polynomial is fitted to the empirical distribution of the total amount of interactions per column, and the first mode of this distribution is used to define the exclusion threshold value below which columns will be removed. This removal should not affect more than 5% of the columns in the matrix.

After the column removal, the remaining bins are further normalized to remove local genomic biases (e.g., to correct for the genomic regions with higher mappability and/or PCR amplification). The normalization procedure implemented in TADbit is a modification of the iterative correction method of HiClib [31]. In the scheme implemented in HiClib, the raw counts for column *i* and row *j* are iteratively corrected until the sum of counts in all the rows converges towards a given value. In TADbit, the iterative correction is stopped when the difference between the columns counts is less than 10% or when a maximum number of iterations (set to 10) is reached, which accelerates the process.

### Comparison of interaction matrices

Once normalized, the Hi-C contact matrices can be compared to estimate their degree of similarity. For this purpose, TADbit implements plotting functions (Supplementary Text) that allow visualizing the interaction matrices as squared heat-maps, in which the two axes represent the genomic coordinates of the analyzed region and the color intensity is proportional to the interaction counts in log2. Besides matrix visualization, TADbit implements two comparison scores: (i) a Spearman rank correlation between bins in two matrices at increasing genomic distances (Fig. 2c) and (ii) a Pearson correlation between the first eigenvectors of each matrix (Fig. 2d). Although both measures aim at identifying whether two matrices are similar or not, they have different properties. The first one is sensitive to the matrix resolution and decays as the genomic distance of the compared bins increases. The second one provides a more global comparison of the matrices and aims at identifying whether the internal correlations in the matrix (detected by its principal eigenvectors) are similar between the compared matrices. In typical Hi-C experiments, one expects the first three eigenvectors to be highly correlated between two similar interaction matrices. If two interaction matrices are similar (that is, with a Spearman rank correlation > 0.2 for all the genomic distances and the Pearson correlation of the first three eigenvectors > 0.7), they may be merged into a single experiment directly within TADbit.

### Genome segmentation into Topologically Associating Domains (TADs)

TADbit analyzes the contact distribution along the genome and subsequently segments it into its constitutive TADs, with each TAD border corresponding to a vertical slice of the Hi-C interaction matrix. TADs can be computed on the interaction matrix from a single experiment or from the matrix resulting from the merge of different experiments. To calculate the position of borders between TADs along a chromosome, TADbit employs a breakpoint detection algorithm that returns the optimal segmentation of the chromosome under BIC-penalized likelihood.

The number of interactions between loci *i* and *j* separated by *Δ* nucleotides is assumed to have a Poisson distribution with parameter *w_ij_ exp(α + β Δ)*, where *α* and *β* are TAD-dependent constants and *w_ij_* is the normalization factor for the cell at coordinates (*i,j*) of the Hi-C contact matrix. Breakpoint detection methods were developed to segment time series in uniform blocks. In the case of Hi-C data, the correspondence with times series is not straightforward because the measured signal is two-dimensional. This issue is resolved by considering that a single observation is the vector of interactions of a locus with all other loci, in other words, an observation is a column of the Hi-C matrix. In this view, a TAD defines a vertical slice of the Hi-C matrix. Each cell of this slice belongs to one of three categories: the contacts between the TAD and all upstream loci, the intra-TAD contacts, and the contacts between the TAD and all downstream loci. From there, the algorithm proceeds in two phases. In the first, the log-likelihood of every slice (defined by a start and end position) is computed. If the slice does not cover exactly one TAD, at least one of the three categories described above will be composite, which will cause a misfit. As a result, the total log-likelihood of this slice will be low. If the slice covers exactly one TAD, all three categories will be uniform and the log-likelihood of the slice will be high. The search for the optimal decomposition of the Hi-C matrix is carried out by a dynamic programming algorithm based on the following property: if *L_k_(s,e)* denotes the log-likelihood of the optimal segmentation of the slice (s,e) into *k* sub-slices, then

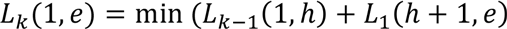

where the minimum is taken over all the values of *h*. This formula allows computing the optimal segmentation recursively.

To assign a border score or strength value, the likelihood of each TAD border in the optimal segmentation is penalized by a value equal to the expected gain in log-likelihood for adding a TAD border after the optimum is reached, and the dynamic programming segmentation is restarted. The whole process is carried out 10 times, and each time a border is on the optimal segmentation, it is penalized by this constant. The strength of a TAD border is the number of times it was included in the optimal segmentation, and it thus ranges from 1 to 10. TAD borders with a score greater than 5 will be considered “robust”, meaning that they are reproducible among different runs; conversely, TAD borders with a score lower than 5 will be considered “weak”, and are likely to be undetectable in replicates or at other resolutions.

### Alignment of TAD boundaries

TAD borders are conserved across different cell types and even across species, indicating that topological domains may play an important role in the organization of chromatin in metazoan genomes [3]. To assess whether TAD borders are conserved throughout different experiments, we implemented a multiple-experiment border alignment algorithm. Starting from different border definitions of the same genomic region, TADbit aligns each TAD to a consensus TAD list, either using the classic Needleman-Wunsch algorithm [36] or using a method based on reciprocal closest boundaries *(bd)*. In the latter method, the boundary *bd1* will be aligned to its closest boundary *bd2* if and only if *bd1* is the closest boundary of *bd2*. Finally, to assess its statistical significance, the resulting alignment is compared to a randomized set of borders obtained by shuffling the TADs, or by randomly picking TADs from a distribution of TAD lengths built from the original alignment.

### Three-dimensional (3D) modeling of genomic domains

In TADbit, the three-dimensional (3D) models of selected genomic domains are generated by transforming the input 3C-based interaction maps into a set of spatial restraints that are later satisfied using the Integrative Modeling Platform (IMP) [37], as previously described [12]. Briefly, 3D modeling with TADbit is based on three main steps: (i) the chromatin domain to be modeled is represented by a set of particles, one per experimental bin of the interaction matrix; (ii) the input interaction data is z-scored and translated into spatial restraints between pairs of particles; and (iii) using Simulated Annealing and Monte Carlo sampling, the imposed restraints are satisfied by building an ensemble of structures whose contact maps correlate as much as possible with the input interaction matrix.

### Structural clustering of the resulting 3D models

To assess the structural similarity of the generated models, TADbit first structurally aligns them using a pair-wise rigid-body superposition that minimizes the RMSD between the superimposed conformations [38]. Then, the *S_mirror_* score of a given pair of models *i* and *j* is computed as:

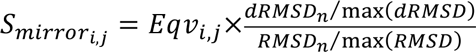

Where *Eqv_ij_* is the number of equivalent positions between two superimposed structures within a specific distance cut-off; *dRMSD_n_* is the normalized (*i.e*., the dRMSD divided by the maximal dRMSD in all structural comparisons) distance RMSD between two aligned structures; and *RMSD_n_* is the normalized RMSD between two aligned structures. This results in a comparison matrix, consisting of all-against-all *S_mirror_* scores, which is then used to resolve structural mirrors (conformations with the same IMP objective function that are mirrors of each other). Next, the comparison matrix is input to the Markov Cluster Algorithm (MCL) program [39] for generating unsupervised sets of clusters of structurally related models. Once the clusters are defined, a representative model of each cluster is compared to obtain the final *dRMSD* used to build the dendrograms of structural similarity between clusters (Fig. 4a). The resulting clusters can be then used independently for the analysis of the generated 3D structures, avoiding thus misinterpretations of the structural variability inherent to the population-based nature of 3C-based experiments.

### Structural analysis of the resulting 3D models

We have implemented a series of structural analysis in TADbit to be applied on the generated 3D models (see online documentation and tutorials http://3dgenomes.github.io/TADbit) and outputs several measures to describe the architecture of the model. Here, we detail the implementation of the four measures used to analyze the structural properties of the fly chromatin types: *particle accessibility, particle density, particle interactions*, and *particle angle*.

The *particle accessibility* assesses whether a chromatin locus in the 3D models is accessible to a hypothetical macromolecule (*e.g*., a transcription factory). To compute this measure, a mesh surface around each particle in the model is first generated. Each point of the mesh represents the center of an object of a user-defined radius (75 nm by default). Next, TADbit checks for each of the points of this mesh whether another particle lies within the specified radius. In that case, that specific mesh point is considered buried and may not be accessed by any other particle or macromolecule. Finally, the accessibility of each particle is computed as the percentage of non-buried mesh points around it. It is important to note that when modeling only part of a genome, the non-modeled part is not taken into account in the accessibility measure and thus its result needs to be interpreted with caution.

The *particle density* assesses the amount of DNA base pairs “packed” within the sum of Euclidian distances between a given particle and its *n* immediate neighbors. Thus, the density measure is simply calculated as the number of base pairs per nm of chromatin fiber.

The *particle interactions* assesses whether a particular locus in the 3D models is in spatial proximity with other loci. Therefore, this measure depends on a distance cutoff value that is set depending on the scaling factor internally used in TADbit to relate genomic distances to Euclidian distances.

The *particle angle* measures the angle (in degrees) formed by three consecutive particles in the 3D models. Similarly as for the density measure, the angle can be computed between *n* distant particles.

Each of these statistics can be calculated independently on consecutive particles or smoothed by averaging the measures in windows across a user-defined number of particles.

### Output and visualization of 3D models

Although TADbit includes a simple three-dimensional model viewer using matplotlib [40], it is designed to be compatible with other visualizing tools, including TADkit (http://www.3DGenomes.org/TADkit). Currently, TADbit can generate three output formats for the model 3D coordinates:

- A simple XYZ format storing the 3D Cartesian coordinates of each particle, as well as any description of the model domain that the user might have previously documented through TADbit. This is a *per model* format, which results in an individual file per model.
- A Chimera Marker format (CMM), which can be used as input for the Chimera molecular viewer [41]. This is also a *per model* format.
- A JSON format, designed to be scalable and broadly used, containing a description of the Hi-C experiment, the input restraints used during the modeling, any additional description of the model domain that the user might have previously documented through TADbit, the Cartesian coordinates of all the models and the model clusters, in case they have been previously calculated by TADbit. This is a *per ensemble* format, which results in a single file per experiment. This file format is fully compatible with the 3D genome browser TADkit.

## Acknowledgments

The research leading to these results has received funding from the European Research Council under the European Union,s Seventh Framework Programme (FP7/2007-2013) / ERC grant agreement 609989, the Spanish Ministry of Economy and Competitiveness (BFU2013-47736-P) and the Human Frontiers Science Program (RGP0044). This work reflects only the authors, views and the Union is not liable for any use that may be made of the information contained therein. We acknowledge support of the Spanish Ministry of Economy and Competitiveness, ‘Centro de Excelencia Severo Ochoa 2013-2017’, SEV-2012-0208 to the CRG.

